# Evidence of interaction between humoral immunity and drug half-life in determining treatment outcome for artemisinin combination therapy in high transmission settings in western Kenya

**DOI:** 10.1101/2020.08.12.248724

**Authors:** Ben Andagalu, Pinyi Lu, Irene Onyango, Elke Bergmann-Leitner, Ruth Wasuna, Geoffrey Odhiambo, Lorna J. Chebon-Bore, Luicer A. Ingasia, Dennis W. Juma, Benjamin Opot, Agnes Cheruiyot, Redemptah Yeda, Charles Okudo, Raphael Okoth, Gladys Chemwor, Joseph Campo, Anders Wallqvist, Hoseah M. Akala, Daniel Ochiel, Bernhards Ogutu, Sidhartha Chaudhury, Edwin Kamau

## Abstract

The role of humoral immunity on the efficacy of artemisinin combination therapy (ACT) has not been investigated, yet naturally acquired immunity is key determinant of antimalarial therapeutic response. We conducted a therapeutic efficacy study in high transmission settings of western Kenya, which showed artesunate-mefloquine (ASMQ) and dihydroartemisinin-piperaquine (DP) were more efficacious than artemether-lumefantrine (AL). To investigate the underlying prophylactic mechanism, we compared a broad range of humoral immune responses in cohort I study participants treated with ASMQ or AL, and applied machine-learning (ML) models using immunoprofile data to analyze individual participants’ treatment outcome. We showed ML models could predict treatment outcome for ASMQ but no AL with high (72-92%) accuracy. Simulated PK profiling provided evidence demonstrating specific humoral immunity confers protection in the presence of sub-therapeutic residual mefloquine concentration. We concluded patient humoral immunity and partner drug interact to provide long prophylactic effect of ASMQ.

## Introduction

Therapeutic efficacy studies (TESs) are used to monitor efficacy of antimalarial drugs including assessment of clinical and parasitological outcome for artemisinin-based combination therapies (ACTs), the first-line treatment for uncomplicated *Plasmodium falciparum* malaria in most endemic countries (1). TESs conducted at regular intervals in the same location can be used for the detection in the decline of drug efficacy over time. Key indicators monitored during ACTs TESs for the treatment of *P falciparum* malaria include proportion of patients who are parasitemic on day 3, and treatment failure by days 28 or 42 (1). It is important, however, to carefully interpret TESs results because they can be influenced by factors originating from the host, the parasite and/or the drugs. Naturally acquired immunity is a key determinant of antimalarial therapeutic response (2), which is highly influenced by transmission intensity (3), and age of the patient (4). It is plausible that TES data may be misinterpreted as population immunity shifts with decline in transmission due to improved malaria control and elimination efforts (5). Pharmacokinetics (PK) and pharmacodynamics of artemisinin derivatives and partner drugs in ACTs are also important when interpreting TES data.

Artemether-lumefantrine (AL) is the most widely used ACTs in sub-Saharan Africa (sSA), followed by artesunate-amodiaquine (ASAQ) (6). A study that investigated clinical determinants of early parasitological response to ACTs in African patients found that risks of persistent parasitemia on the first and the second day were higher in patients treated with AL compared to those treated with dihydroartemisinin-piperaquine (DP) and ASAQ (7). However, on the third day, the difference was not apparent. Artesunate-mefloquine (ASMQ) has been extensively used in Asia and Latin America but not in sSA because of the availability of other more affordable ACTs (8), concerns for mefloquine resistance seen in Southeast Asia (SEA) (9), and side effects such as excessive vomiting in children (10). In addition, AL and ASAQ remain highly efficacious (6), and there has not been conclusive evidence or motivation for the need to introduce ASMQ (11). Nonetheless, the World Health Organization (WHO) has recommended ASMQ be reconsidered for the treatment of uncomplicated malaria in sSA (1).

Even though ACTs remain highly efficacious in sSA (6), a recent study conducting genetic analysis to evaluate *P falciparum* parasite genomes across Africa identified recent signature of selection on chromosome 12 with candidate resistance loci against artemisinin derivatives evident in Ghana, and Malawi (12). This calls for continued surveillance on the efficacy of ACTs across Africa while refining approaches and tools used because of the heterogeneous nature of transmission across the continent. One such strategy that has been proposed is the use of multiple or alternating first-line ACTs in order to extend lifespan of these drugs (13). This will require availability and deployment of additional ACTs. To this end, we conducted a TES in western Kenya, a high transmission region where we compared efficacy of ASMQ and DP to AL, the first-line treatment for uncomplicated malaria in Kenya. Protein microarray with more than a thousand *P falciparum* antigens were used to identify malaria antigen-specific antibody responses that were significantly different between study participants with respect to ASMQ and AL treatment outcomes on days 28 and 42. Machine learning and statistical modeling were used to assess the degree to which the humoral immunity to malaria could be used to make individualized predictions of treatment outcomes when ASMQ and AL are used.

## Materials and Methods

### Study design and participants

This was a randomized, open-label, two-cohort trial, each with two arms conducted in western Kenya, a high transmission, holoendemic region. Cohort I study was conducted between June 2013 and November 2014, and assessed ASMQ and AL, while cohort II study was conducted between December 2014 and July 2015, and assessed DP and AL. Patients aged between 6 months and 65 years presenting with uncomplicated malaria at the local outpatient referral hospital were recruited into the study. Potential study participants were identified from malaria rapid diagnostic test (mRDT) positive patients. Screening procedures were undertaken only after obtaining informed consent/assent from the participants, parents or legally authorized representatives. In order to be considered for enrollment, the study participants had to demonstrate willingness and ability to comply with the study activities for the duration of the study, including willingness to become in-patient for the first three days following enrolment. Consort figure (Figure 1) provide additional details. Further, study details are described in the Supplementary material.

**Figure 1.**
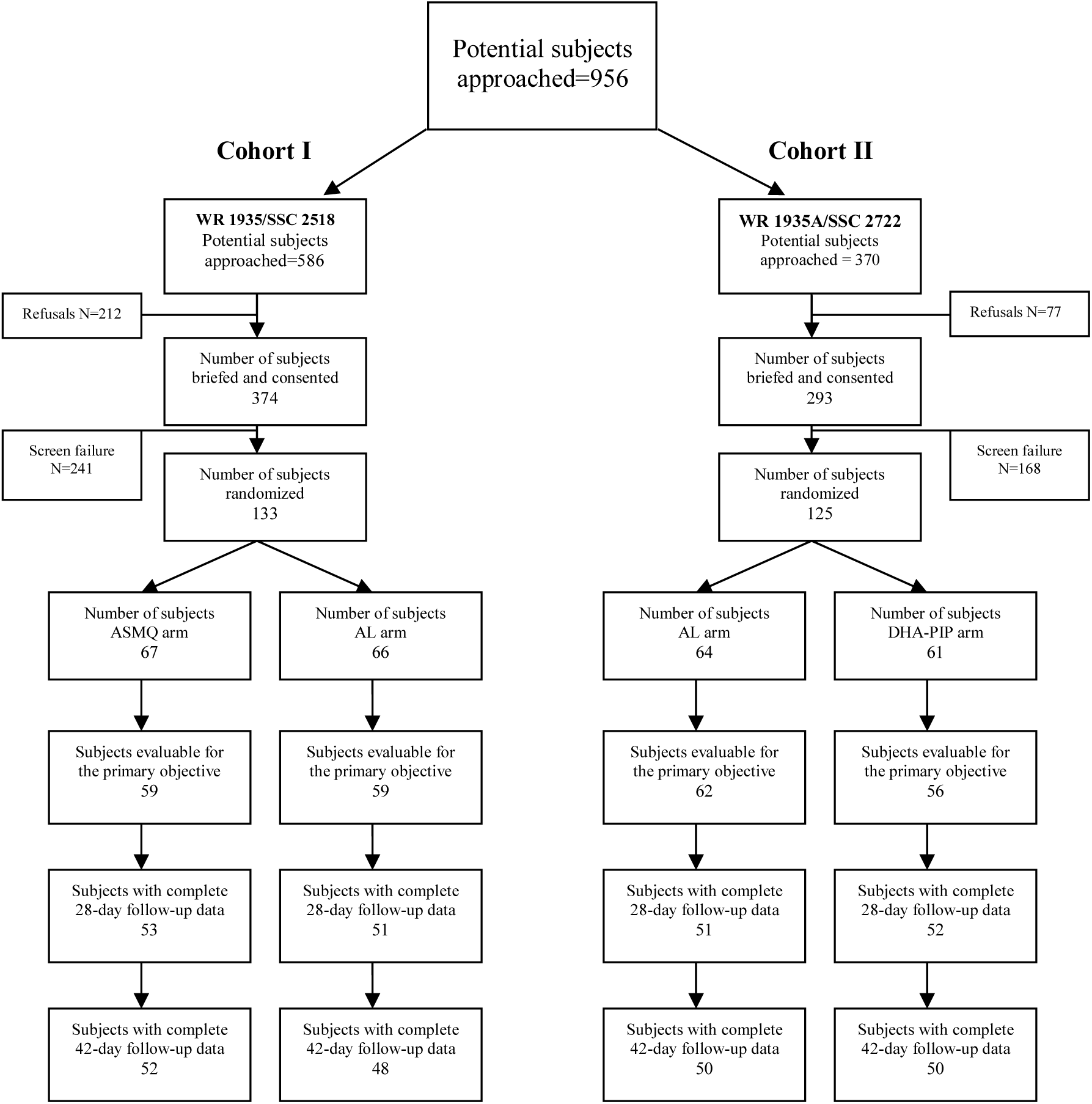
Consort Figure. Consort figure. Detail information on recruitment and execution of the study.

### Study procedures

Enrolled participants were given one of the malaria treatments that were being investigated based on the randomization schemes: DP (Duo-cotecxin® - Holly Cotec Pharmaceuticals, Beijing, China), ASMQ (available as separate artesunate and mefloquine tablets obtained from the WHO, Geneva, Switzerland) or AL (Coartem® - Novartis Pharma Ag, Basel, Switzerland). DP and AL were administered orally at the standard dosage per the drug package insert according to the pre-defined weight bands for each of the drugs. ASMQ was administered orally in a staggered fashion: artesunate was administered on its own at hour 0, 24 and 48 at the standard dose of 4mg/kg; mefloquine was then administered at hour 72 (at 15mg/kg) and hour 96 (at 10 mg/kg), thus totaling to 25mg/kg.

During the treatment phase of the studies, blood samples were collected at hours 0, 4, 8, 12, 18, 24, and thereafter, 6 hourly until 2 consecutive negative smears for malaria were obtained. Upon completion of study treatment, participants were followed up weekly from day 7 through day 42. During these follow-up visits, blood samples were collected for malaria testing. Participants found to have malaria during the weekly follow-up visits were treated as per the National Guidelines for the Diagnosis, Treatment and Prevention of Malaria in Kenya.

### Study outcomes

WHO definitions for treatment outcomes in malaria drug efficacy studies were used (1). The primary study endpoint was the time to parasitemia clearance from day 0 after treatment initiation. Parasite clearance rates were calculated using the Worldwide Antimalarial Resistance Network (WWARN), Parasite Clearance Estimator (PCE) tool located at http://www.wwarn.org/toolkit/data-management/parasite-clearance-estimator). Log transformed parasite density was plotted against time in hours to generate the slope half-life which is defined as the time needed for parasitemia to be reduced by half.

### Laboratory procedures

Malaria microscopy was performed on thick and thin smears prepared using standardized procedures. Quantification was done based upon a complete blood count of a sample drawn on the same day as the smear. The complete blood counts were performed using validated Coulter AcT 5diff hematology analyzers (Beckman Coulter, Pasadena, CA). The findings of two (or three in case of discrepancies) independent expert microscopists were considered. *In vitro* drug sensitivity testing was conducted on day 0 pre-treatment samples as well as on samples collected from participants who had reappearance of parasites on follow-up visits utilizing the malaria SYBR Green technique as previously described (14). Details on how molecular tests were performed can be found in the Supplementary Material.

### Protein microarrays and Ab profiles

A protein microarray containing a total of 1087 *P falciparum* antigens were produced by Antigen Discovery Inc. (ADI, Irvine, CA, USA) from the 3D7 proteome as previously described (15). A detailed description of production as well as execution of the experiments are provided in Supplementary material. Blood samples collected at enrollment before the initiation of treatment were used in microarray experiments. Due to resource limitations and personnel attrition that are not uncommon in resource-challenged study environments such as those of sSA (where the study was conducted), immunoprofiling was carried out for cohort I only (ASMQ and AL arms). Subsequent bioinformatics, data analysis and modeling were performed only for cohort I, which the challenges and shortcoming have been highlighted in the discussion.

### Bioinformatics, data analysis and modeling

Detailed bioinformatics, data analysis and modeling methods can be found in the Supplementary material. Briefly, to identify antibody signal intensities that differed with respect to different treatment outcome, univariate analysis was conducted for each antibody signal in the immunoprofile. ASMQ and AL study arms were analyzed separately. Within each arm, participants were further classified as treatment success or treatment failure based on non-PCR-corrected Adequate Clinical and Parasitological Response (nPC-ACPR) on day 28 and day 42 per World Health Organization (WHO) definition and guidance for the treatment of malaria (1). Each antibody signal was compared between treatment success (nPC-ACPR = 1) and treatment failure (nPC-ACPR = 0). Random forest and logistic regression were applied to build machine learning models using all antibody signals to predict individual participants’ treatment outcome (nPC-ACPR). To evaluate the predictive accuracy of random forest models, cross-validation was utilized, where data samples were subsampled by up-sampling. This aggregation for training and prediction performance was evaluated on data samples that were not used in training. Models’ performance was expressed as both a percentage of correctly predicted outcomes with a Cohen’s kappa value, and as the area under the curve of the receiver operating characteristic (AUCROC). Cohen’s kappa statistic is a measure that can handle imbalanced class problems. A kappa value > 0.4 can indicate that the classifier is performing better than a classifier that guesses at random according to the frequency of each class. To assess the statistical significance of the models and check the overfitting that might occur in the machine learning process, AUCROC-based permutation tests were carried out. Relative importance scores of each antibody signal were calculated using random forest models. Principal component analysis (PCA) was applied to all the antibody signals with relative importance scores > 50. Antibody signal intensities of participants in the ASMQ arm were plotted using principal component (PC)1 and PC2. We constructed one-compartment PK models for artemether, artesunate, lumefantrine and mefloquine, respectively, using the *linpk* R package. The values of PK parameters, including bioavailable fraction, central clearance, central volume, and first-order absorption rate, were obtained from a previous study (16). All statistical analyses were performed using the R *stats* package and STATA version 13 (StataCorp) while machine learning was carried out using the R *caret* package.

## Results

### ASMQ and DP outperforms AL in parasite clearance

A total of 956 (586 in cohort I and 370 in cohort II) potential study participants were recruited, 236 were enrolled in the study, 118 in each cohort. Out of 236 study participants, 200 (84.7%) completed 42-day follow-up, 100 from each cohort. The consort figure provides a full breakdown of study participants recruitment and execution (Figure 1). There were no notable differences in the baseline characteristics of the enrolled participants (Supplementary Table 1). There were no cases of early treatment failure observed in both study cohorts (Supplementary Figure 1), and all study participants achieved 100% PCR-corrected ACPR (PC-ACPR) rates at day 28 and day 42 (Table 1). There were no significant differences in the parasite clearance half-lives between the study arms regardless of the treatment used (Table 2). The maximum parasite clearance slope half-lives observed for both cohorts were 4.2 (AL arm in cohort I) and 4.3 (AL arm in cohort II) hours (Table 2), which falls within the WHO recommended cut-off of five hours for suspected artemisinin resistance (1). However, PC50 and PC99 data which is defined as the time taken for the initial parasite density to fall by 50% or 99% clearly showed ASMQ and DP outperformed AL by clearing parasites much faster (Table 2). In cohort I, ASMQ cleared 99% of the parasites on average 5.3 hours fasters than AL, with maximum time of 25.6 hours compared to 33.0 hours in AL. In cohort II, DP cleared 99% of the parasite on average 3.0 hours faster than AL, with maximum time of 27.9 hours compared to 30.5 hours in AL. Further, participants who received ASMQ and DP achieved better nPC-ACPR at day 28 and day 42 compared to those who received AL (Table 1), with significant difference present in cohort I for ASMQ vs AL on day 28 (p = 0.042) but not on day 42 (p = 0.280), and in cohort II, significant difference was present for DP vs AL on day 28 (p = 0.001) and on day 42 (p = 0.008).

**Table 1:** Rates of adequate clinical and parasitological response (ACPR) with and without PCR corrections

|  | ASMQ | AL | Difference %<br>(95% CI) | DP | AL | Difference %<br>(95% CI) |
| --- | --- | --- | --- | --- | --- | --- |
| **Day 42** | **n = 52** | **n = 48** |  | **n = 50** | **n = 50** |  |
| **PC-ACPR** | 52(100%) | 48(100%) |  | 50(100%) | 50(100%) |  |
| **nPC-ACPR** | 30(57.7%) | 21(43.8%) | -13.9%(-33.4 to 5.5) | 39(78.0%) | 21(42.0%) | -36%(-53.8 to -18.1) |
| **Day 28** | **n = 53** | **n = 51** |  | **n = 53** | **n = 51** |  |
| **PC-ACPR** | 53(100%) | 51(100%) |  | 53(100%) | 51(100%) |  |
| **nPC-ACPR** | 45(84.9%) | 32(62.8%) | -22.2%(-38.6 to -5.8) | 50(96.2%) | 36(70.6%) | -25.6%(-39.1 to -12.0) |

**Table 2:** Parasite clearance rates. Data shows parasite slope half-lives and parasite clearance rates for cohort I (ASMQ and AL-I) and cohort II (DP and AL-II) in hours.

| Parameters | ASMQ | AL-I | DP | AL-II |
| --- | --- | --- | --- | --- |
| **Total Analyzed for T<sub>1/2</sub>** | **58** | **50** | **57** | **53** |
| **Slope half-life Median(IQR)** | 2.3(1.8-2.7) | 2.5(2.2 – 3.0) | 2.2(1.9-2.5) | 2.3(2.0-2.9) |
| **Slope half-life Mean(range)** | 2.2(0.98-3.6) | 2.6(1.5-4.2) | 2.2(1.2-3.6) | 2.4(1.4-4.3) |
| **PC50 Median(IQR)** | 4.0(2.6 – 6.0) | 7.4(4.8 – 9.6) | 4.1(3.2 – 6.0) | 6.7 (4.4 – 8.8) |
| **PC50 Mean(range)** | 4.2(0.28-11.1) | 7.4(0.5-15.3) | 4.6(0.27-11.3) | 6.5(0.24-11.6) |
| **PC99 Median(IQR)** | 17.0(13.4 – 19.2) | 21.5(18.3 – 24.8) | 16.8(14.6 – 19.5) | 20.1(17.5 – 22.1) |
| **PC99 Mean(range)** | 16.6(6.5-25.6) | 21.9(10.0-33.0) | 17.0(7.3-27.9) | 20.0(9.1-30.5) |

### Study participants experienced minimal adverse events

Table 3 lists some of the adverse events that occurred. With exception of anemia, tinea capitis and respiratory tract infections, frequency of adverse events were extremely low, mild and self-limiting. Of note, we did not have a single participant vomit, show mental or neurological side-effects, including those in ASMQ arm.

**Table 3:** Adverse events. Some of the adverse events that occurred in study participants. None of the study participant vomited, all adverse events were mild, at low frequency and resolve spontaneously.

|  | ASMQ | AL | DP | AL |
| --- | --- | --- | --- | --- |
| Number of patients | n = 59 | n = 59 | n = 62 | n = 56 |
| Anemia | 15(25.4%) | 9(15.3%) | 7(11.3%) | 10(17.9%) |
| Diarrhea | 0 | 0 | 1(1.6%) | 0 |
| Abdominal pain | 1(1.7%) | 0 | 1(1.6%) | 0 |
| Gastroenteritis | 1(1.7%) | 0 | 2(3.2%) | 0 |
| Tinea capitis | 5(8.5%) | 3(5.1%) | 4(6.5%) | 4(7.1%) |
| Respiratory tract infection | 3(5.1%) | 15(25.4%) | 12(19.4%) | 9(16.1%) |
| Rash | 2(3.4%) | 1(1.7%) | 0 | 3(5.4%) |
| Dizziness | 1(1.7%) | 0 | 0 | 0 |
| Fatigue | 0 | 1(1.7%) | 0 | 0 |
| Headache | 1(1.7%) | 1(1.7%) | 5(8.1%) | 1(1.8%) |
| Fever | 0 | 0 | 1(1.6%) | 0 |
| Nausea | 1(1.7%) | 0 | 0 | 0 |

### In vitro and molecular parasite analyses

*In vitro* susceptibility testing to AL component drugs (artemether and lumefantrine) was successfully performed in some of the parasite isolates (Supplementary Figure 2). The parasite isolates IC_50_ values for both drugs remained unchanged and they were within range of previous published IC_50_ values of parasite isolates from this region (17). K13 mutations were present in some of the parasite isolates, with N632K being the most prevalent in cohort I and L663I in cohort II. However, none of the K13 mutations identified as markers of artemisinin resistance in SEA were present in these samples. Interestingly, the type, amount and proportion of K13 mutations were significantly different in samples from the two cohort studies, indicating that these mutations are transient and not related to ACT drug pressure. None of the K13 mutations were associated with increased parasite slope half-lives (Supplementary Figure 3).

We investigated the polymorphisms in *pfcrt* (K76) and *pfmdr1* (N86, 184F and D1246), and *pfmdr1* copy numbers which are associated with AL selection in sSA parasites (17). In cohort I study, 97% of the samples carried wildtype *pfcrt* K76 and *pfmdr1* N86, 82% carried wildtype D1246, and 41% carried mutant *pfmdr1* 184F. In cohort II, 100% of the samples carried wildtype *pfcrt* K76, 99% carried wildtype *pfmdr1* N86, 89% carried wildtype D1246, and 43% carried mutant *pfmdr1* 184F. These frequencies are similar to what has previously been reported for parasites from this region (17). None of these mutations or haplotypes were associated with increased parasite slope half-lives (Supplementary Figure 4). There was no difference in *pfmdr1* copy number variation between the two cohort studies, with 90% and 92% of the parasites carrying a single copy of the gene in cohort I and II, respectively.

### Higher humoral immunity confers better nPC-ACPR outcome in ASMQ arm

Once we had established that the variances in the response to drug treatment were not due to differences in parasite genetic diversity profiles, we sought to determine whether distinct immunoprofiles in participants treated with the different drug combinations impacted the clinical and parasitological outcome in cohort I. Serological immunoprofiling using Pf1000 protein microarrays was successfully performed for 91 (46 in ASMQ and 45 in AL) of the 104 participants who completed day 28 follow-up, and 87 (45 in ASMQ and 42 in AL) of 100 participants who completed day 42. We carried out univariate analyses to compare antibody signals to the 1087 antigens contained in the microarrays between participants who achieved nPC-ACPR vs those who did not in ASMQ and AL arms, on day 28 and day 42. Significant nPC-ACPR associated differences were present in the ASMQ arm (Figure 2A and B), but not in the AL arm (Figure 2C and D). In the ASMQ arm, antibody responses to 277 antigens with *P* < 0.05 and adjusted *P* < 0.1 (Figure 2A) were significantly different between participants that showed nPC-ACPR on day 28 compared to those who did not. On day 42, significant differences were observed for antibody responses to 10 antigens with *P* < 0.05 and adjusted *P* < 0.1 (Figure 2B). The right-skewed pattern in the volcano plots (Figure 2A and B) indicates that participants maintaining nPC-ACPR in the ASMQ arm had higher humoral immunity to *P falciparum* antigens compared those who did not. The specific antigens that correspond to these antibody responses are listed in Supplementary Table 2, ranked by corresponding Benjamini-Hochberg adjusted *P* values. When we investigated the stages of the *P falciparum* lifecycle for the 277 antigens associated with ASMQ treatment outcome, we found they were overrepresented in sporozoites and trophozoites stages (Supplementary Table 3).

**Figure 2.**
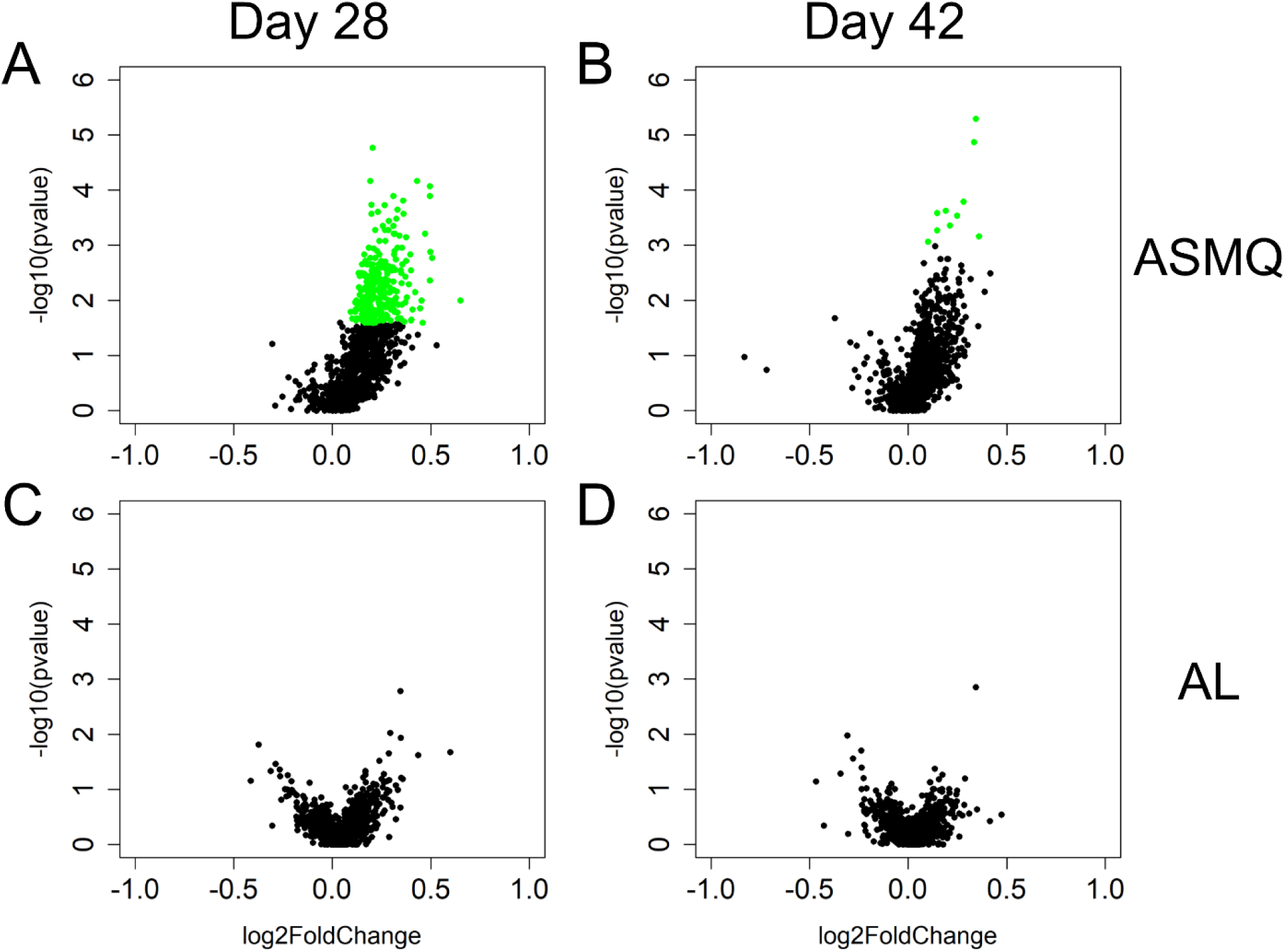
nPC-ACPR-associated differences in subjects’ humoral immunity to malaria in the ASMQ and AL arms. Univariate analyses were applied to identify differences in subjects’ humoral immunity associated with ASMQ outcomes on day 28 (A) and day 42 (B), and with AL outcomes on day 28 (C) and day 42 (D), respectively. Volcano plots were used to present analysis results. The x-axis is log2 ratio of malaria antigen-specific antibody signals of subjects presenting nPC-ACPR to those of subjects not presenting nPC-ACPR. The y-axis is *P* values based on – log10. The green dots represent the antibody responses with *P* < 0.05 and Benjamini-Hochberg adjusted *P* < 0.1.

To determine whether distinct immunoprofiles are associated with parasitological outcome, all microarray and clinical data were integrated and a PCA performed. In the ASMQ arm, participants clustered separately based on their treatment outcome, showing clear systematic differences in immune responses (Figure 3A and 3B). However, there was no separation in the AL arm (Figure 3C and 3D).

**Figure 3.**
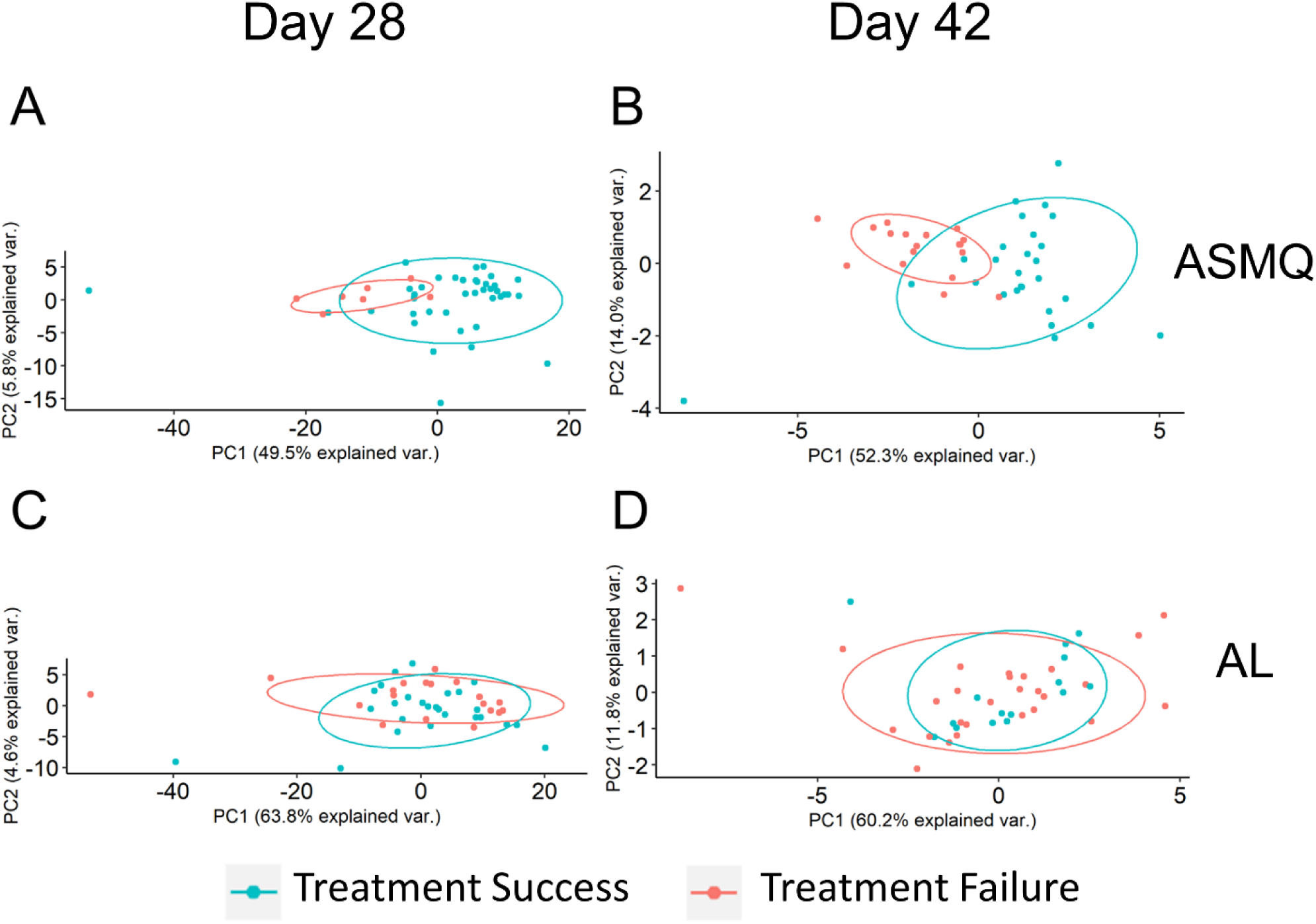
Principal Component Analysis (PCA) plots of treatment outcome-specific differences. (A) and (B). PCA used antibody signals with adjusted P < 0.1 to visualize treatment outcome-specific differences in the ASMQ arm, which were identified by univariate analyses applied to identify differences in subjects’ humoral immunity associated with ASMQ outcomes on day 28 and day 42, respectively. (C) and (D). PCA used the same antibody signals as (A) and (B) to visualize treatment outcome-specific differences in the AL arm.

### Antibody responses associated with nPC-ACPR are age dependent

To investigate the magnitude of immune responses with age, we compared antibody signal intensities of the study participants in different age groups in the ASMQ arm. Overall, signal intensities of younger participants were lower than those of older participants. Normalizing the mean signal intensities to *P falciparum* proteins from participants at varying ages against the mean intensities for the oldest subject group revealed that the magnitude of antibody responses to these antigens increased with age (Figure 4). Among antibody responses associated with nPC-ACPR on day 28, children (participants < 12 years) showed approximately half the magnitude of antibody responses as older participants (≥ 12 years), as indicated by the slope. This effect was more pronounced in antibody responses associated with nPC-ACPR on day 42, where children showed approximately a third of the magnitude of responses as older participants. These findings suggest that antibody responses associated with nPC-ACPR are acquired over time through repeated exposures as children age into adults, thus corroborating previous studies (4, 18, 19).

**Figure 4.**
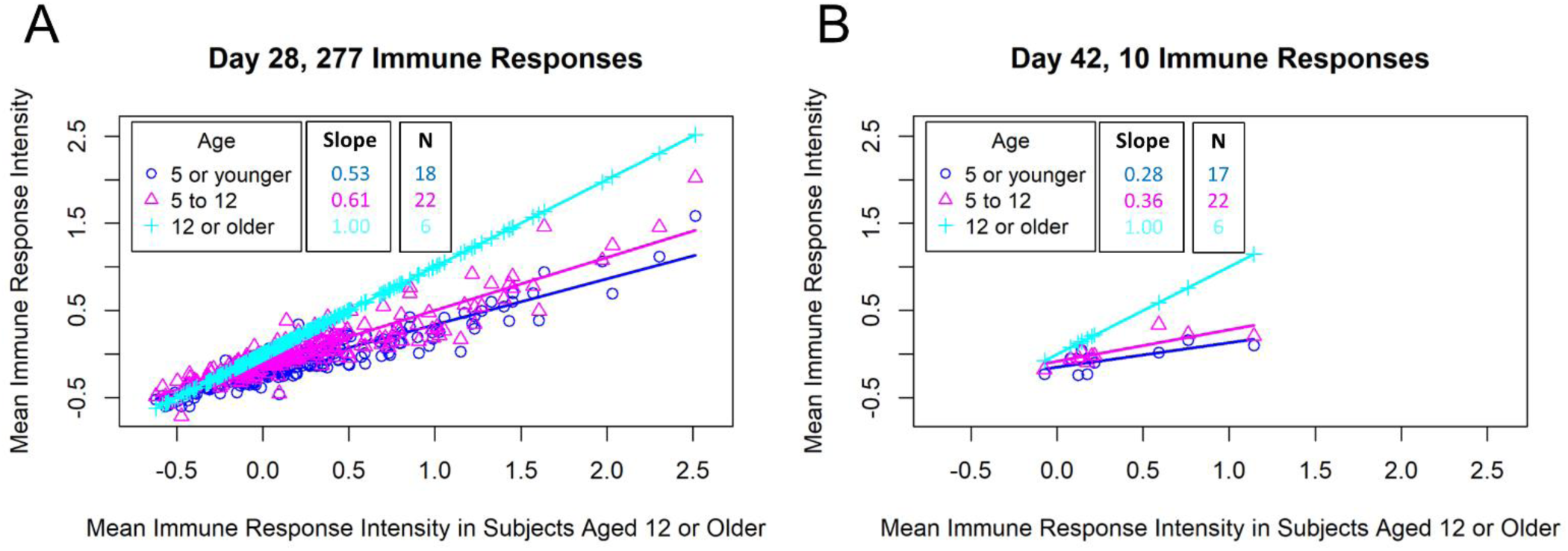
Comparison of malaria antigen-specific antibody responses of subjects in different age groups. The mean antigen-specific antibody signal intensities to *P falciparum* proteins (identified through univariate analyses) from subjects in varying ages (y-axis) were plotted against the mean intensities for the oldest subject group (x-axis). (A) Scatterplot of mean antibody signal intensities of 277 *P falciparum* proteins associated with ASMQ outcomes on day 28.

### Machine learning can predict treatment outcome for ASMQ using humoral immunity data

Machine learning methods were used to assess the degree to which humoral immunity to malaria could be used to make individualized predictions of treatment outcome. Data presented here was generated using random forest model but was confirmed by running logistic regression models that provided similar findings. For the ASMQ arm, 100 random forest models were built for predicting treatment outcomes using antibody signals that were significantly different between participants that showed nPC-ACPR compared to those who did not. Models built for predicting treatment outcomes on day 28 achieved 85% accuracy (Kappa: 0.40) with an average AUCROC of 0.85 (Figure 5), and on day 42, the accuracy was 72% (Kappa: 0.43) with an average AUCROC of 0.83 (Figure 5). Models built on top two antibody signals (rifin [PF3D7_0900200_e2s1_19] and GTP binding protein [PF3D7_1411600_383]) achieved 92% accuracy for both day 28 and 42 (Supplementary Figure 5). Randomly shuffled nPC-ACPR outcomes across participants to remove any possible link between humoral immunity and outcome were used as negative control for the machine-learning analysis to test for overfitting. The average AUCROC for using the randomly shuffled data for day 28 and 42 in ASMQ arm were 0.54 and 0.53 (Figure 5C and D), indicating that the day 28 and day 42 ASMQ models were not over-fitted. In the AL arm, machine learning models could not predict nPC-ACPR outcome on day 28 or day 42, achieving accuracies of 53% (Kappa: 0.02) and 61% (Kappa: 0.05), respectively, with an average AUCROC of 0.51, indicating an accuracy no better than random chance (Figure 6).

**Figure 5.**
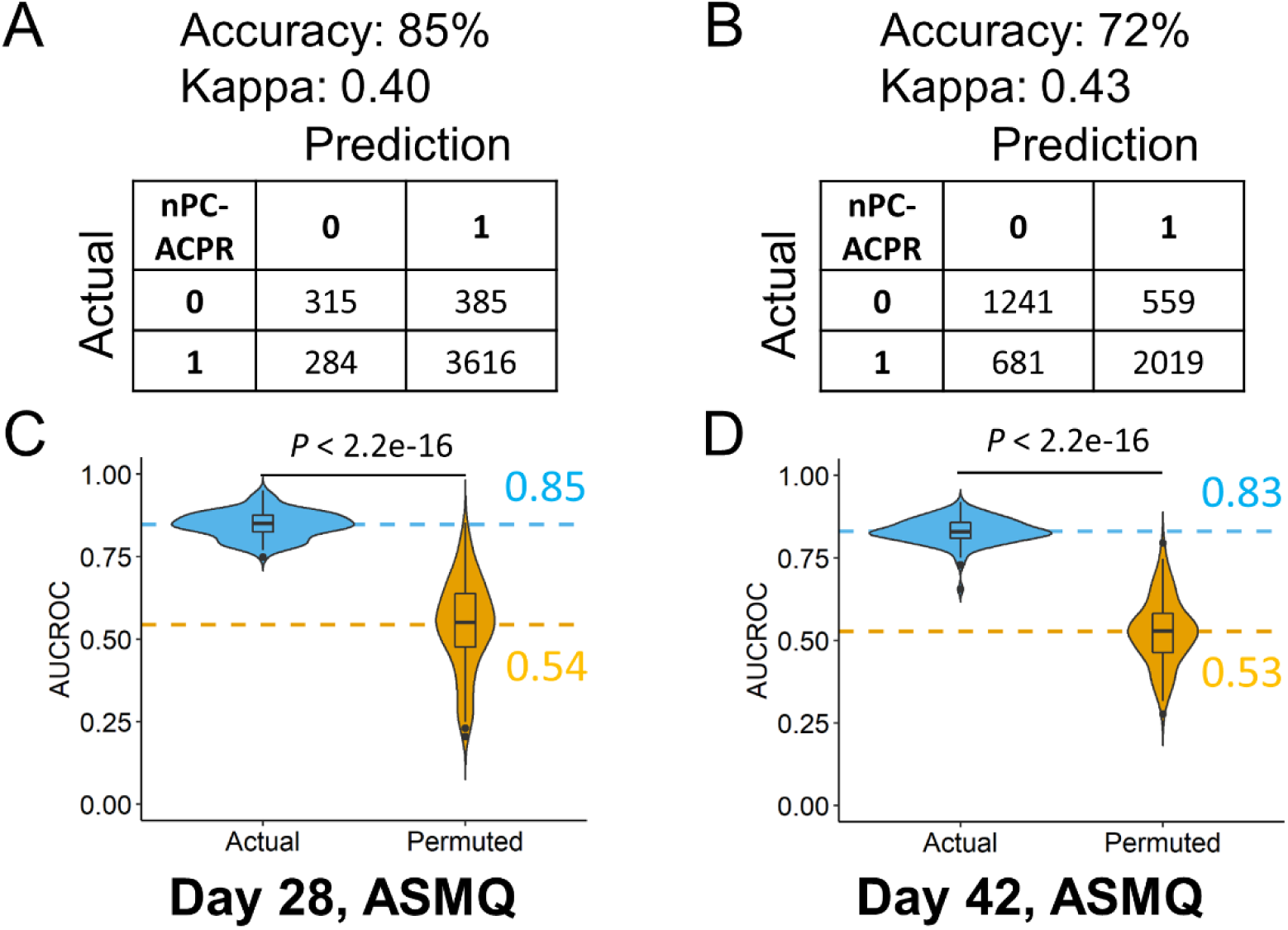
Performance evaluation of random forest models predicting ASMQ outcomes on day 28 and day 42. (A) and (B). Prediction accuracy, kappa, and confusion matrices. The rows of confusion matrices represent the predicted treatment outcomes, whereas the columns indicate the actual treatment outcomes. (C) and (D). Comparison of AUCROC values from 100 repetitions of 100 times repeated 5-fold cross-validation using actual (blue) versus permutated (yellow) nPC-ACPR labels. Dashed line represented the mean AUCROC values. Significance is determined using Mann-Whitney-Wilcoxon test.

**Figure 6.**
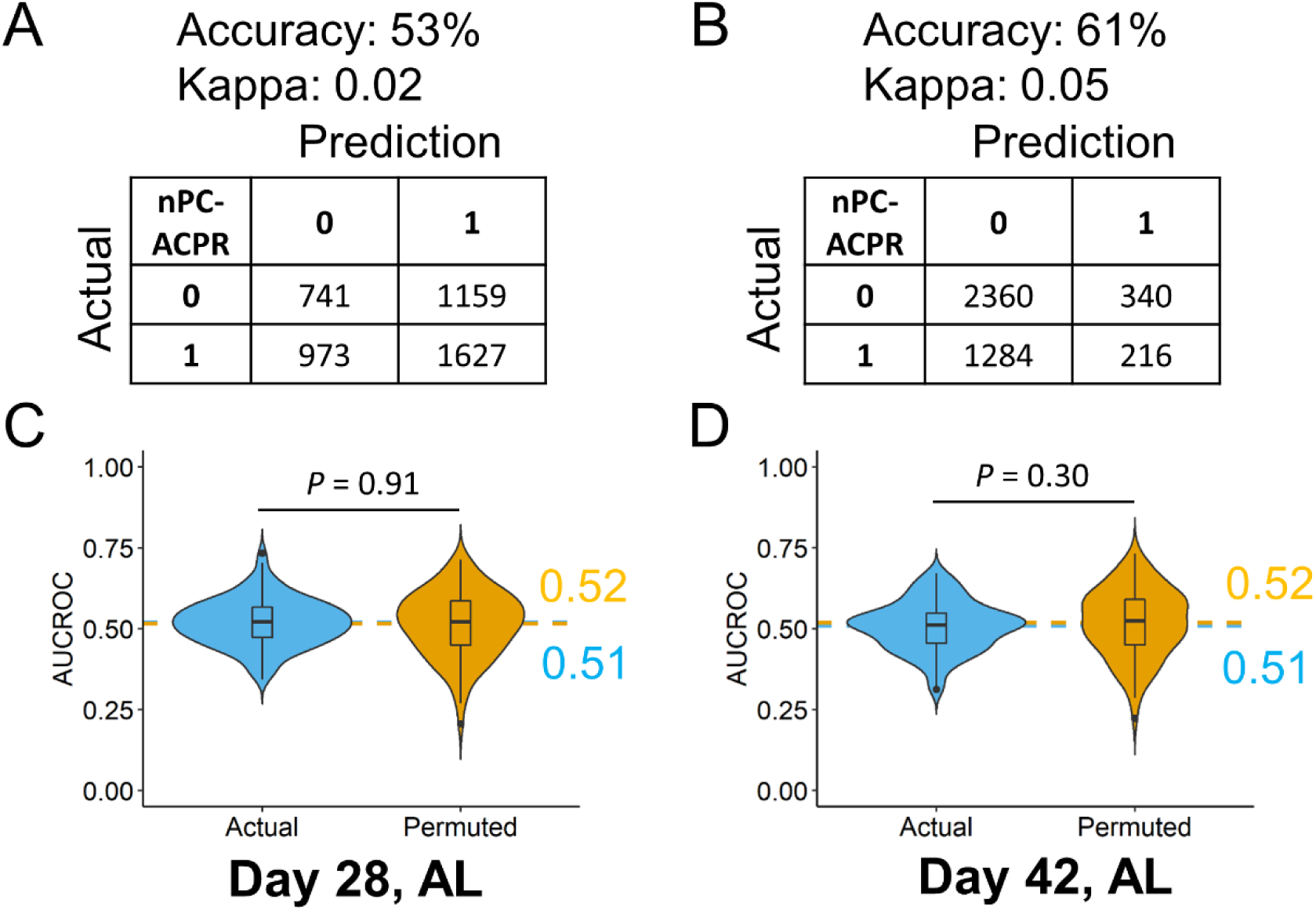
Performance evaluation of random forest models predicting AL outcomes on day 28 and day 42. (A) and (B). Prediction accuracy, kappa, and confusion matrices. The rows of confusion matrices represent the predicted treatment outcomes, whereas the columns indicate the actual treatment outcomes. (C) and (D). Comparison of AUCROC values from 100 repetitions of 100 times repeated 5-fold cross-validation using actual (blue) versus permutated (yellow) nPC-ACPR labels. Dashed line represented the mean AUCROC values. Significance is determined using Mann-Whitney-Wilcoxon test.

### Simulation of ASMQ and AL PK profiles

To explore how humoral immunity and treatment interact, we simulated PK profile of ASMQ and AL out to day 28 and day 42. The PK profiles of both regimens show that the concentration of artemisinin derivative drugs clears within 6 days post-treatment. In the AL regimen, lumefantrine is cleared by day 11 post-treatment, therefore is unlikely to have a major impact on day 28 and day 42 outcomes (Figure 7A and Supplementary Figure 6). By comparison, in the ASMQ regimen, mefloquine has a much longer apparent half-life, with concentrations relative to peak concentration of 30.9% at day 28, and 3.3% at day 42 (Figure 7A), representing 3- and 30-fold reductions from the peak concentrations.

**Figure 7.**
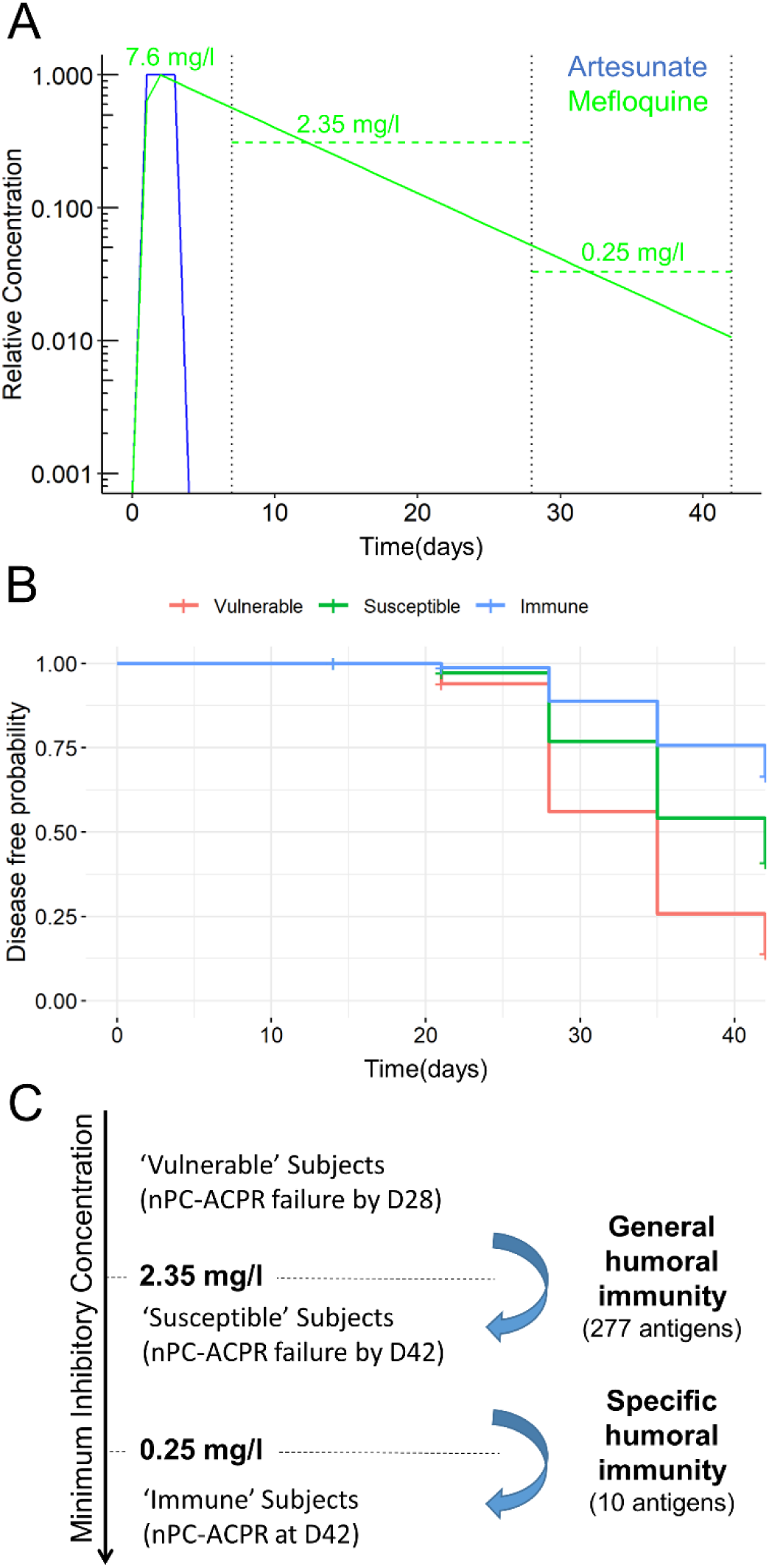
PK profile of ASMQ and estimation of MIC of mefloquine. A) Simulated PK profile of ASMQ for artesunate (blue) and mefloquine (green). Estimated peak concentration, and average concentration between day 7 and day 28, and day 28 and day 42 are shown for mefloquine are labelled. B) Estimated disease-free probability for all ASMQ subjects classified as ‘susceptible’, ‘intermediate’, and ‘immune’ based on humoral immunity using the day 28 and day 42 predictive models. C) Relationship between humoral immunity and MIC of mefloquine based on ACPR outcome, univariate analysis, and simulated PK profiles.

These results led us to formulate the hypothesis that humoral immunity augments the efficacy of mefloquine when it is at sub-therapeutic concentrations. The minimum inhibitory concentration (MIC) of the drug required to be effective is likely to vary between individuals. Given that all participants showed nPC-ACPR at day 7, for participants that showed nPC-ACPR failure by day 28, the drug concentration necessarily fell below individual’s MIC between day 7 and day 28. Based on our simulated PK profiles, the average mefloquine concentration during that time span was 2.35 mg/l. Likewise, for participants that showed nPC-ACPR at day 28, but nPC-ACPR failure by day 42, the mefloquine concentration fell below their MIC within the time span of day 28 and day 42, during which an average concentration was estimated to be 0.25 mg/l. Therefore, based on our estimates, ‘vulnerable’ participants who had nPC-ACPR failure by day 28 have an average MIC > 2.35 mg/l, while immune participants who show nPC-ACPR even out to day 42 have an average MIC of < 0.25 mg/l (Figure 7A). In addition, Cox regression analysis shows that there is a strong association (*P* < 0.01) between the time to malaria re-infection and classes of participants identified by machine learning models (Figure 7B).

## Discussion

Our study provides crucial findings on the efficacy of ACTs in treatment of uncomplicated *P falciparum* in high transmission settings. To the best of our knowledge, this is the first study to demonstrate divergent impact of serological immune profiles on treatment outcomes based on ACT treatment. Using machine learning, we identified *P falciparum* antigens that are highly predictive of successful ASMQ treatment outcome. Our modeling data suggest that at sub-therapeutic concentrations, mefloquine act synergistically with malaria-specific antibody responses to provide extended protection against clinical and parasitological failure. In sSA, there is a need to deploy additional ACTs or new class of antimalarial drugs to avoid development of resistance to the current first-line treatments. By identifying specific antigens associated with, and predictive of treatment outcome for specific antimalarial drugs, our data support the notion of smart deployment of new ACTs and other antimalarial drugs, where decisions are informed by individual and population immune profiles, and strategically considered for each region or a country.

This study was conducted in Kombewa district hospital under stringent, supervised conditions. Treatments were administered precisely in terms of dosage, timing and meal requirements in an in-patient hospital setting. The high efficacy of all the ACTs tested in this study can be attributed to high transmission rates resulting in high immune status, and in part to the rigor with which the study was conducted. This is especially true for ASMQ which was dispensed as a loose, non-fixed dose combination, which has been shown to elevate risk of poor treatment outcome (20). We showed ASMQ and DP outperformed AL on day 28, which became much more apparent on day 42, corroborating earlier studies (7, 11). This has previously been attributed to the long half-lives of mefloquine and piperaquine, which are thought to provide 4-6 weeks prophylaxis after treatment compared to 3-4 days for lumefantrine (21). Using PK modeling, we estimated mefloquine concentrations to be at 30% of peak concentration on day 28, while lumefantrine was completely cleared by 14 days, confirming the persistence of mefloquine, and its role in the apparent post-treatment prophylaxis observed in the ASMQ cohort. With changing and highly heterogeneous transmission in sSA, this study highlights the importance of considering transmission intensity when evaluating therapeutic efficacy of current, or new antimalarial drugs including combination therapies.

Host immunity has previously been shown to be an important determinant of treatment outcome in *P falciparum* malaria infections (2), with the magnitude of immunological response increasing with age (4). In this study, we have shown the interaction between humoral immunity and residual mefloquine concentration is important in providing protection and predicting treatment outcome. This is supported by the following observations: First, if humoral immunity alone was sufficient for nPC-ACPR out to day 28 and day 42, then immunity would have predicted protection in AL arm as well; second, if residual mefloquine concentration alone was sufficient to determine nPC-ACPR outcome, then humoral immunity wouldn’t have predicted outcome in ASMQ; and third, immunity is likely the only explanation for differences in nPC-ACPR based on age, as age-specific differences in PK profiles of AL and ASMQ have not been reported. Machine learning identified three classes of patients (vulnerable, susceptible, and immune) in the ASMQ arm based on immune data (Figure 7B). Our findings suggest that general humoral immunity to a wide range of malaria antigens is sufficient to provide protection in the presence of residual mefloquine concentrations out to day 28, while specific immunity to a handful of select antigens is necessary to provide protection in very low residual mefloquine concentrations out to day 42. Using PK modeling to estimate MICs, we found that this specific immunity results in a dramatic 10-fold reduction in mefloquine MICs. Given the highly variable nature of host immunity and its close link to transmission intensity, this study highlights the importance of considering host immunity and transmission intensity when evaluating the efficacy of new or existing antimalarial drugs.

*P falciparum* protein microarray have been used to investigate antibody responses predictive of protection from symptomatic malaria in high (22) and seasonal transmission (15) settings. These studies had few common antibodies that were identified as predictive of protection, some of which were similar to those present in our study (Supplementary Table 2). Notably, antigens considered as vaccine candidates provided conflicting results in these studies, where they were predicted as protective in one study (22) but not in the other (15). In our study, none of the vaccine candidate antigens were predicted to influence treatment outcome. Unlike the previous studies, we computationally integrated antibody profiles with clinical and parasitological outcome, and used machine learning to identify antigens associated with treatment outcomes at different time points.

Reports of mefloquine side-effects including early vomiting, mental and neurological concerns might be contributing to the poor scale-up of ASMQ in Africa (10, 11, 23). However, studies that have reported ASMQ as fixed- or non-fixed-dose combination have shown it is effective and well-tolerated, with minor advance events that seem to vary widely between studies (11, 23). Additionally, reports of elevated mefloquine side effects out of Africa (Nigeria and Malawi) are from monotherapy studies (24, 25), which have not been replicated elsewhere on the continent. It seems dosing and timing of when mefloquine is administered as a combination therapy is critical, which impacts drug efficacy, and the side effects experienced by the patient. Early vomiting is related to age, disease severity and a history of vomiting in the previous 24 hours, which is likely to be influenced by the immune status of the patient. Kuile *et al*., showed that in children ≤ 2 years, vomiting was reduced by 40% when mefloquine dose of 25 mg/kg was split over two days, and by 50% when given on the second day (10). By administering artesunate first, and then mefloquine 24 hours later, this reduced vomiting because the patients had recovered clinically and were more likely to tolerate mefloquine. Further, delaying the dose of mefloquine for 24 hours after artesunate administration increases mefloquine oral bioavailability substantially probably due to rapid clinical improvement (26).

In our study, we administered three doses of artesunate in the first 48 hours, and then mefloquine at hours 72 and 96, at 15 mg/kg and 10 mg/kg, respectively. This dosing scheme eliminated vomiting in the ASMQ arm and dramatically reduced reported side-effects. As a fixed-dose or non-fixed-dose combination therapy, ASMQ is given over a three-day period once or twice daily (23, 27-29). Given the importance of dispensing medication as fixed-dose for improved compliance, we propose creation of a fixed-dose ASMQ combination that delivers mefloquine after the first 24 or 48 hours (preferably) to allow ample time for clinical recovery of the patient, which is likely to improve treatment outcome. Further, since malaria symptoms and mefloquine side effects are difficult to distinguish, studies that investigate the short and long-term side effects in healthy immune and non-immune African populations (children and adults), as therapeutic or prophylaxis are warranted.

This study had several limitations. Firstly, we did not perform microarray analysis of samples from cohort II study, and neither did we perform PK simulation using individual data from our study participants. This was mainly due to time and funding shortfalls. In the future, it will be important to repeat these analyses using other ACTs especially DP due to its high efficacy and the long prophylactic life-span of piperaquine. However, we are confident in the results of the PK analysis because data used was from a study conducted in western Kenya, the same patient population as those in our study.

In conclusion, we have demonstrated that data integration, machine learning, and modeling provide a comprehensive approach capturing the underlying complexity of malaria control in sSA. Further, we have shown ASMQ is a highly effective drug in this setting, with treatment outcome dependent on patient immune status. By changing the dosing schedule, we have shown side effects associated with ASMQ can be drastically reduced. Genetic analysis revealed parasite population in this location remain highly susceptible to ACTs, and there are no genetic signatures associated with mefloquine resistance, making ASMQ an appropriate choice of possible first-line treatment in western Kenya, a region which account for most malaria transmission in the country.

## Acknowledgments

We thank Dr Veronica Manduku, KEMRI Center for Clinical Research; LTC Claire A Cornelius, Dr Douglas Shaffer, Directors, USAMRD-A/K, and Dr. Steve Munga, KEMRI Center for Global Health Research, for supporting this study and giving their permission to publish these data. We also thank all clinical staff at Kombewa District Hospital for their assistance. But most importantly, we would like to pour our heartfelt gratitude to the study participants whom sacrifice a lot to participate in our studies for the betterment of the community, thank you!

## Disclaimer

The opinions or assertions contained herein are the private views of the author, and are not to be construed as official, or as reflecting true views of the Department of the Army, the U.S. Department of Defense, or The Henry M. Jackson Foundation for Advancement of Military Medicine, Inc (HJF). The investigators have adhered to the policies for protection of human subjects as prescribed in AR 70–25. Material has been reviewed and approved for public release with unlimited distribution by the Walter Reed Army Institute of Research and HJF.

Financial support: Funding for this study was provided by the Armed Forces Health Surveillance Branch (AFHSB) and its Global Emerging Infections Surveillance (GEIS) Section, Grant P0126_13_KY. The study sponsor had no role in study design; in the collection, analysis, and interpretation of data; in the writing of the report; and in the decision to submit the paper for publication.

## Supplementary Data

### Methods

#### Study design and participants

The study was conducted in Kombewa, a high transmission, holoendemic region located in western Kenya where malaria transmission occurs throughout the year with two seasonal peaks around April to August and October to December coinciding with long and short rainy seasons, respectively. This study was a randomized, open-label two-cohort trial, each with two arms designed to capture the different transmission seasons. The cohort I study was conducted between June 2013 and November 2014, assessing AL and ASMQ, and the cohort II was conducted between December 2014 and July 2015, assessing AL and DP. Patients aged between 6 months and 65 years inclusive presenting with uncomplicated malaria at the outpatient clinic at the local referral hospital located in Kombewa were recruited into the study. Potential study participants were identified from malaria rapid diagnostic test positive patients. Screening procedures were undertaken only after obtaining informed consent/assent from the participants, parents or legally authorized representative. The study participants had to provide willingness and ability to comply with the study protocol for the duration of the study including willingness to remain in hospital for three days. Screening procedures included obtaining a blood sample for malaria parasite molecular identification, genetic analysis, quantification by microscopy and complete blood count. Participants who were enrolled were admitted at the local referral hospital for approximately three days for treatment administration and intensive blood sample collection.

#### Sample size determination

The primary objective for both cohorts was to determine parasitological clearance rates by microscopy within the first 72-hour period after first ACT dose in patients with uncomplicated Pf malaria. At least 59 participants per arm (considering 20% loss to follow-up) had to be recruited into each arm. This sample size would give a power of 34.6% to detect a difference in the parasite clearance half-life (ΔK) of 0.01 between the study arms using a pairwise t-test. A precision of 20% (ΔK=0.02) would result in the following differences in parasite clearance time: K=0.1: half-life=3.01 hours, 90% clearance time =10.0 hours K=0.1+20%: half-life=2.51 hours, 90% clearance time= 8.33 hours

#### Study procedures

Enrolled participants were given one of the treatments for malaria that were investigated in the studies as per the randomization schemes: AL (Coartem® - Novartis Pharma Ag), ASMQ (available as separate artesunate and mefloquine tablets obtained from the World Health Organization) or DP (Duo-cotecxin® - Holly Cotec Pharmaceuticals). AL and DP were administered orally at the standard dosage as per the drug package insert according to the pre-defined weight bands for each of the drugs. ASMQ was administered orally in a staggered fashion: artesunate was administered on its own at hour 0, 24 and 48 at the standard dose of 4mg/kg; mefloquine was then administered at hour 72 (at 15mg/kg) and hour 96 (at 10 mg/kg), thus totaling to 25mg/kg. Treatment was repeated for participants who vomited within 30 minutes of administration. If they vomited a second time, they were withdrawn from the study and offered parenteral treatment as per the National Guidelines for the Diagnosis, Treatment and Prevention of Malaria in Kenya (1).

During the treatment phase of the studies, blood samples for various tests (including microscopy, molecular and genetic testing, hematology and pharmacokinetic assays) were collected at hours 0, 4, 8, 12, 18, 24, and thereafter, 6 hourly until 2 consecutive negative smears for malaria were obtained. Upon completion of study treatment, participants were followed up weekly from day 7 through day 42. During these follow up visits, blood samples were collected for testing as indicated above. Participants found to have malaria during the weekly follow-up visits were treated as per the National Guidelines for the Diagnosis, Treatment and Prevention of Malaria in Kenya (1).

#### Study outcomes

The World Health Organization (WHO) definitions for treatment outcomes in malaria drug efficacy studies were used (2). The primary study endpoint was the time to parasitemia clearance from day 0 after treatment initiation. Parasite clearance rates were calculated using the Worldwide Antimalarial Resistance Network (WWARN) tool for Parasite Clearance Estimator (PCE; located at http://www.wwarn.org/toolkit/data-management/parasite-clearance-estimator). Log transformed parasite density was plotted against time in hours to generate the slope half-life which is defined as the time needed for parasitemia to be reduced by half.

#### Laboratory procedures

##### Microscopy

Malaria microscopy was done on one thick and one thin blood smear prepared from each sample collected using WHO standardized procedures (2). Parasitological assessments of thick and thin blood films were performed independently by two expert microscopists. Quantification was done based upon a complete blood count of a sample drawn on the same day as the smear. The complete blood counts were performed using validated Coulter AcT 5diff hematology analyzers (Beckman Coulter). The findings of 2 (or 3 in case of discrepancies) independent expert microscopists were considered.

##### In vitro analysis

*In vitro* drug sensitivity testing was conducted on all day 0 pre-treatment samples as well as on samples collected from participants who had reappearance of parasites on follow-up visits using the malaria SYBR Green technique as previously described (3, 4).

##### Molecular detection assays

Detection of *Plasmodium* was performed by real-time qPCR as previously described using QuantiFast PCR Master Mix (5). Mutations in the Kelch 13 (K13)-propeller domain were analyzed by sequencing as previously described (6). Single Nucleotide Polymorphisms (SNPs) associated with drug resistance including *Plasmodium falciparum* (Pf) chloroquine resistance gene (*pfcrt* 76) and Pf multidrug resistance gene (*pfmdr1*; 86, 184, and 1246), were screened using PCR-based single base extension on Sequenom MassARRAY system (Agena Bioscience, San Diego, CA, USA) as previously described (7). Data analysis was performed using CLC Main workbench 7 with reference to the PF3D7_1343700 control sequence retrieved from PlasmoDB (www.plasmodb.org).

##### Recrudescence and re-infection assays

To distinguish between recrudescence and re-infection if treatment failure occurs, Pf obtained from blood samples collected during follow-up visits were compared to those taken at baseline using polymorphisms within three genes: merozoite surface protein-1 (MSP1), merozoite surface protein-2 (MSP2), and glutamate-rich protein (GLURP). MSP1 was typed in a nested PCR reaction for allelic typing for MAD20, for K1 and for R033 family alleles (8). MSP2 allelic type was determined using a nested PCR assay targeting the FC27 and 3D7 alleles (9). Genotyping using the GLURP gene was done using hemi-nested PCR using as previously described (9). An outcome was defined as recrudescence if a subsequent sample contained identical alleles or a subset of the alleles present in the first sample. An outcome was defined as re-infection if a subsequent sample contained only new alleles. If a subsequent sample contained alleles present in the first sample and new alleles, the outcome initially was considered indeterminate.

#### Protein microarrays and Ab profiles

A protein microarray containing a total of 1087 Pf antigens were developed by Antigen Discovery Inc. (ADI, Irvine, CA, USA) from the 3D7 proteome as previously described (10). Briefly, proteins were expressed from a clone library generated at ADI in an *E. coli* cell-free in vitro transcription and translation (“IVTT”) system (Rapid Translation System, 5 Prime, Gaithersburg, MD, USA). Proteins were spotted on nitrocellulose-coated glass AVID microarray slides (Grace Bio-Labs, Inc., Bend, OR, USA) using an Omni Grid Accent robotic microarray printer (Digilabs, Inc., Marlborough, MA, USA). Microarrays were packed in padded slide boxes protected from light and shipped to USAMRD-A in Kisumu, western Kenya where the assay was performed as follows. The microarrays were blocked using 200 µl Blocking buffer (Whatman Inc, Sanford, ME) per well for 30-60 minutes on a slow platform rocker. Serum samples were diluted 1:100 in protein array blocking buffer and pre-absorbed in 10% DH5α *E. coli* lysate to block anti-*E. coli* antibodies that could bind to IVTT components on the spots and mask protein-specific responses. The slides were incubated over-night at 4°C with gentle agitation. Slides were washed using wash buffer and then incubated for one hour at room temperature with Biotin-conjugated goat anti-human IgG Fcγ fragment specific secondary antibody (Jackson ImmunoResearch, West Grove, PA, USA) diluted in 1:1000 blocking buffer. Tertiary detection was done by diluting Streptavidin-conjugated SureLight P-3 (Columbia Biosciences, Frederick, MD, USA) in 1:200 blocking buffer and then incubating for one hour at room temperature covered with aluminum foil in the dark. The slides were washed with wash buffer, then in ultra-pure water and air-dried by centrifugation for 5 minutes at 500 rpm. At this point, microarray slides were stored in padded slide boxes protected from light and return shipped to ADI for scanning and quantification of signals. Slides were scanned on a GenePix 4300A High-Resolution Microarray Scanner (Molecular Devices, Sunnyvale, CA, USA), and raw spot and local background fluorescence intensities, spot annotations and sample phenotypes were imported and merged in R, in which all subsequent procedures were performed. Slide images were visually assessed for contaminations and evidence of drying during the assay and shipment, and where effects were present, spot signals were censored. Foreground spot intensities were adjusted by local background by subtraction, and negative values were converted to one. All foreground values were transformed using the base two logarithm. The dataset was normalized to remove systematic effects by subtracting the median signal intensity of the IVTT control spots on each array for each sample. With the normalized data, a value of 0.0 means that the intensity is no different than the background, and a value of 1.0 indicates a doubling with respect to background.

#### Bioinformatics, data analysis and modeling

##### Univariate analysis

To identify antibody (Ab) signal intensities that differed with respect to different treatment outcomes, univariate analysis was conducted for each Ab signal for cohort 1 study. AL and ASMQ study arms were analyzed separately. Within each arm, participants were further classified as treatment success or treatment failure based on non-PCR-corrected ACPR (nPC-ACPR) on day 28 and 42. Each Ab signal was compared between treatment success (nPC-ACPR = 1) and treatment failure (nPC-ACPR = 0) participants (nPC-ACPR day 28 = 1 vs. nPC-ACPR day 28 = 0, nPC-ACPR day 42 = 1 vs. nPC-ACPR day 42 = 0). Firstly, Shapiro-Wilks tests were carried out to determine if the to-be-compared Ab signal intensities were normally distributed in both treatment success and treatment failure groups. If both were normally distributed (*P* < 0.05 by the Shapiro-Wilks test), Student’s t-tests were applied between treatment success and failure groups. Otherwise, Wilcoxon signed-rank tests were performed. After *P* values were calculated for each antigen (Ag) signal in the data set, the Benjamini-Hochberg (BH) adjusted *P* values were calculated to correct for false discovery rate. Differences of Ab signal intensities were considered significant at *P* < 0.05 and BH adjusted *P* < 0.1. In addition, differences in Ab signal intensities were compared across age groups (< 5; ≥ 5 to < 12; and ≥ 12 y old).

##### Enrichment analysis

Life cycle stages of Pf Ags on the microarray were determined using previously published mass spectrometry data obtained from PlasmoDB (www.plasmodb.org). Evidence of expression at the Sporozoite, Merozoite, Trophozoites, and Gametocyte stages were obtained from a comparative proteomics study on Pf clone 3D7 (11). Stage expression information of liver stage antigens were inferred from mass spectrometry evidence of orthologues in *P. yoelii* (12). Enrichment analysis was carried out using hypergeometric tests (phyper function within the stats package in R) that were set at the 0.05 level for type I error.

##### Prediction of outcome using machine learning

Random Forest (RF) was applied to build machine learning models using all Ab signals to predict participants’ nPC-ACPR. Model trainings and parameter tunings were carried out using repeated 5-fold cross-validation, subsampling the data set by 5-fold and resampling 100 times. The hyperparameter, mtry (number of variables randomly sampled as candidates at each split), was adjusted to identify the optimal out-of-bag error, an unbiased estimate of the generalization error. To evaluate the predictive accuracy of the RF modeling approach, cross-validation was utilized, where data samples were subsampled by up-sampling. This aggregation for training and prediction performance was evaluated on those observations that were not used in training. To assess the statistical significance of the random forest models and check the overfitting that might occur in the machine learning process, AUCROC-based permutation tests were carried out. In permutation tests, the outcome labels of the training data were shuffled randomly. Each random forest model was rebuilt using the data with permuted labels repeatedly for 100 times. AUCROC was computed to evaluate prediction performance of permutation models. Based on AUCROC of permutation models, null distributions for AUC were also estimated. Logistic regression (LR) models were built to confirm the findings of RF models. The hyperparameters, alpha (the elasticnet mixing parameter) and lambda (the regularization parameter), were tuned using repeated 5-fold cross-validation to identify the optimal AUCROC.

##### Principal component analysis

Principal component analysis (PCA) was applied to all the Ab signals with relative importance scores > 50. Ab signal intensities of participants in the ASMQ arm were plotted using principal component (PC)1 and PC2 that captured the most and second most variations from the data, which could determine if participants could be clustered by their treatment outcomes. Treatment outcomes on day 28 and day 42 were analyzed separately. *Pharmacokinetic (PK) modeling:* We constructed one-compartment PK models for artemether, artesunate, lumefantrine and mefloquine, respectively, using the R *linpk* package. The values of PK parameters, including bioavailable fraction, central clearance, central volume, and first-order absorption rate, were obtained from previous studies (13, 14). PK simulations were performed to produce PK concentration-time profiles of each medication for 42 days post treatment. Artesunate and mefloquine were compared with respect to their relative concentrations (normalized by peak concentrations)-time profiles, so were artemether and lumefantrine.

##### Cox regression analysis

A Cox proportional-hazards model was built to investigate the association between the time to malaria reinfection and the classes of participants identified by machine learning modeling, using the R *survival* package. The time to malaria reinfection was determined by the malaria testing, weekly from day 7 through day 42. All statistical analyses were performed using the R *stats* package and machine learning were carried out using the R *caret* package.

---

**Supplementary Table 1:** Baseline characteristics of cohort I (ASMQ and AL-I) and cohort II (DP and AL-II) study participants

|  | AL - I<br>N=59 | ASMQ<br>N=59 | AL - II<br>N=62 | DP<br>N=56 | Overall<br>N=236 |
| --- | --- | --- | --- | --- | --- |
| Age – mean years (SD) | 6.7 (7.1) | 8.9 (10.1) | 7.0(3.8) | 7.3 (3.6) | 7.5 (6.7) |
| Age – median years (range) | 5 (1 – 50) | 6 (1 – 51) | 6 (1 – 22) | 7 (2 – 19) | 6 (1 – 51) |
| Sex – female N(%) | 34 (57.6%) | 35 (59.3%) | 31 (50%) | 27 (48.2%) | 127 (53.8%) |
| Mean weight kg (SD) | 24.0 (15.7) | 25.0 (15.1) | 22.2 (10.0) | 22.1 (8.7) | 23.3 (12.7) |
| Weight range | 10.6 – 90.4 | 11.0 – 76.6 | 11.0 – 69.4 | 11.4 – 64.2 | 10.6 – 90.4 |
| Temperature – mean °C (range) | 38.0 (36.0 – 40.1) | 37.5 (36.1 – 40.0) | 37.7 (36.0 – 40.1) | 37.5 (35.9 – 39.6) | 37.7 (35.9 – 40.1) |
| Mean Hb g/dl (range) | 10.6 (7.0 – 15.1) | 10.8 (7.2 – 15.8) | 11.0 (6.8 – 15.3) | 11.0 (6.8 – 15.3) | 10.9 (6.6 – 15.8) |
| Asexual Parasite density – geometric mean per µL (95% CI) | 38759 (26194 – 57351) | 32789 (21907 – 49078) | 36928 (25438 – 53607) | 38652 (24505 – 60964) | 36709 (30080 – 44799) |
| Gametocyte carriage N(%) | 51(86.4%) | 52 (88.1%) | 46 (74.2%) | 49 (87.5%) | 198 (83.9%) |

**Supplementary Table 3:** Percentage of *P falciparum* protein features with mass spectrometry evidence for expression during each of the various parasite lifecycle stages.

| Parasite Stage | Percent of *P falciparum* proteins targeted by 277 selected Ab immune responses, Day 28 (p-value) | Percent of *P falciparum* proteins targeted by 10 selected Ab immune responses, Day 42 (p-value) | Percent of 1,087 *P falciparum* proteins on microarrays |
| --- | --- | --- | --- |
| Liver | 14.9 (0.13) | 10.0 (0.63) | 12.8 |
| Sporozoites | 43.5 (0.03) | 30.0 (0.42) | 38.6 |
| Merozoites | 26.1 (0.41) | 30.0 (0.53) | 26.8 |
| Trophozoites | 31.2 (0.48) | 60.0 (0.05) | 30.8 |
| Gametocytes | 37.7 (0.16) | 30.0 (0.51) | 35.1 |

